# Adolescent intermittent ethanol exposure induces sex-specific and time-dependent changes in affective behaviors and metabolomic profiles

**DOI:** 10.1101/2025.04.29.651256

**Authors:** Mariah J. Shobande, Anjali Kumari, Michael Pearson, Janae A. Baker, Nzia I. Hall, Renee C. Waters, Chloe E. Emehel, Dashear Hill, Myla E. Fowlkes, Tiffany Dean, Reginald Cannady, Bo Wang, Antoniette M. Maldonado-Devincci

**Affiliations:** Department of Chemical, Biological and Bioengineering, College of Engineering, North Carolina Agricultural and Technical State University, Greensboro, North Carolina 27411, U.S.A; Department of Biology, College of Science and Technology, North Carolina Agricultural and Technical State University, Greensboro, North Carolina 27411, U.S.A; Department of Social Work and Sociology, Hairston College of Health and Human Sciences, North Carolina Agricultural and Technical State University, Greensboro, NC 27411; Department of Psychology, College of Health and Human Sciences, North Carolina Agricultural and Technical State University, Greensboro, North Carolina 27411, U.S.A; University of North Carolina at Chapel Hill School of Medicine, North Carolina, 27516, U.S.A; Department of Psychology, Princeton Neuroscience Institute, Princeton University, Princeton, New Jersey, 08540, U.S.A; Department of Chemistry and Chemical Engineering, Florida Institute of Technology, Melbourne, FL 32901-6975; Department of Biomedical and Chemical Engineering and Sciences, Florida Institute of Technology, Melbourne, FL 32901-6975

**Keywords:** Adolescence, Sex Differences, AIE, Mouse, Dependence, Alcohol, Metabolomics, Amino Acids, Serum, Liver, Fecal Samples

## Abstract

Given the adverse effects of adolescent binge alcohol consumption in humans, the present work explores the lasting behavioral and metabolomic impacts following adolescent binge ethanol exposure. Here we determine short- and long-term changes in affective behaviors and metabolomic profiles in male and female adolescent binge ethanol-exposed mice. Male and female C57BL/6J mice were exposed to adolescent intermittent ethanol (AIE) as a model of binge ethanol exposure using intermittent vapor inhalation from postnatal day (PND) 28-42. Mice were tested in a battery of behavioral tests and metabolomic analyses were conducted following short-term and long-term withdrawal from AIE exposure. Mice were tested for affective behaviors using the open field test (OFT), the light/dark test (LDT), and the tail suspension test (TST) one week following AIE exposure from PND 49-53 and again from PND 91-95. Serum samples were collected on PND 43, corresponding to 24 hours after the last exposure, fecal samples were collected during each OFT, and liver samples were collected approximately 80 days after AIE exposure for metabolomic analysis. We show modest incubation of behavioral differences in anxiety-like behavior in males after adolescent binge ethanol exposure; an effect that was absent in female mice. In contrast, metabolomic differences in male mice that were more pronounced shortly after adolescent binge ethanol exposure waned as time progressed since last exposure. Male mice appear to be more susceptible to the persistent changes in adolescent binge ethanol exposure that varies over time. It is possible that short-term metabolomic changes may predict long-term behavioral changes in affective behaviors.

**Contribution to the field statement:** Adolescent binge alcohol exposure causes long-lasting changes in behavior, however the underlying biology mediating these changes have not readily been assessed in a sex-specific manner. Metabolomic assays are a powerful tool that can be used as biomarkers to determine changes in biological pathways that are linked to specific behavioral phenotypes. Here we explored the relationship between adolescent binge alcohol exposure, changes in anxiety-like behavior, and metabolomic profiles in male and female C57BL/6J mice. We find robust changes in metabolomic profiles following short-term withdrawal in male mice exposed to binge alcohol during adolescence. We also see modest changes in anxiety-like behavior following long-term withdrawal. Although female mice did not show any robust changes in anxiety-like behavior nor global changes in metabolomic profiles, in both sexes we do observe persistent changes in some amino acids, which may serve as specific biomarkers associated with changes in alcohol-induced behavioral changes. These data add to the field by conducting a novel longitudinal study following male and female mice from adolescence to early adulthood and measuring physiological biomarkers (metabolomics) coupled with behavioral changes following binge ethanol exposure.

## 1. Introduction

The phenomenon of adolescents engaging in high-risk behaviors, including alcohol drinking, is quite common. Specifically, in 2019 among 16-17 year-old teens, 17.8% of boys and 20.8% of girls reported using alcohol in the past month (SAMHSA 2020). While adolescents drink less frequently than adults, when they do drink, binge drinking is the most common pattern of consumption (Windle 2016), with 10.2% of boys and 11.2% of girls reporting binge drinking in the 2019 National Survey on Drug Use and Health (SAMHSA 2020). Over the past several years, sex differences in binge drinking rates in teens have narrowed, and teenage girls are drinking similar or more alcohol compared to teenage boys (SAMHSA 2020). This early profile in alcohol drinking rates during adolescence sets the stage for a higher percentage of adult women to develop health-related complications faster than men, despite lower overall lifetime alcohol consumption (Fama, Le Berre, and Sullivan 2020; White 2020).

Heavy alcohol exposure can alter physical, cognitive, social, emotional, behavioral, and neural development during adolescence (Hiller-Sturmhöfel and Spear 2018; Spear and Swartzwelder 2014; Thorpe et al. 2020; Marco et al. 2017). Studies in animal models, mostly in rats, show persistent effects of adolescent intermittent ethanol (AIE) exposure on some affective-like behaviors including ethanol drinking, anxiety-like behavior, impulsivity, behavioral flexibility, memory, sleep, and social anxiety (Crews et al. 2019; Dannenhoffer et al. 2018; Pandey et al. 2015; Robinson et al. 2021; Sakharkar et al. 2019; K. L. Healey et al. 2022). Thus, the data from clinical, and some basic research studies, support the conclusion that binge-like ethanol exposure during adolescence creates a propensity toward affective dysregulation (Dannenhoffer et al. 2018; Gilpin 2012; Pandey et al. 2015; Varlinskaya, Kim, and Spear 2017). However, most research to date investigating AIE effects has focused on alterations in neurobehavioral functioning, and little work in animal models has included analyses of AIE-induced changes in peripheral nervous system functioning.

Sex differences in the acute and long-term effects of adolescent ethanol exposure are increasingly found in clinical and preclinical research. In rats, AIE exposure sex-specifically alters affective behaviors including exploratory and anxiety-like behaviors (Varlinskaya et al. 2020; Aguilar et al. 2023; K. L. Healey et al. 2022), fear conditioning (Chandler, Vaughan, and Gass 2022), and social drinking (Towner et al. 2022) in adulthood. We recently observed sex differences following withdrawal from AIE and voluntary alcohol drinking; where females showed changes in anxiety-like behaviors following short-term and long-term withdrawal; while male AIE-exposed mice showed changes in anxiety-like behavior (Maldonado-Devincci et al. 2022) and stress reactivity (K. Healey et al. 2023) only after long-term withdrawal. Other work shows greater AIE-induced changes in affective behaviors (e.g., novelty-induced hypophagia) in females compared to male C57BL/6J mice (Kasten et al. 2020). We also recently showed that AIE induced long-term increases in Grin2b in the cerebellum following long-term abstinence in both male and female mice (K. Healey et al. 2023). Together, these data indicate that AIE alters later behavioral changes in a sex-specific and withdrawal time-dependent manner, but more work is needed to understand the underlying long-lasting biological changes that mediate these behavioral alterations, including differences in alcohol pharmacokinetics and alcohol metabolism seen between sexes (Kezer, Simonetto, and Shah 2021).

Metabolomics is a useful scientific approach to gathering non- to minimally invasive samples to determine which biochemical processes may be altered in the body due to certain stimuli (Costanzo et al. 2022). Metabolomics can be a powerful tool to complement other ‘omics approaches to indicate the function of an organism and to identify specific biomarkers that may be altered following different environmental perturbations (Harrigan, Maguire, and Boros 2008). This approach is used to determine molecular phenotypes of various diseases and disorders like diabetes, cardiovascular disease, liver disease, changes in affective behavior, and alcohol and substance use disorders (Tilg, Cani, and Mayer 2016; Williams et al. 2014; Mostafa et al. 2017; Ghanbari et al. 2021; Voutilainen and Kärkkäinen 2019; Harrigan, Maguire, and Boros 2008; Harada et al. 2016). In humans, alcohol in dependence alters metabolomic profiles, leaky gut, and is associated with higher scores on affective measures of anxiety and depression (Leclercq et al. 2014, 2024). Alcohol use is also linked to changes in metabolites including amino acids, lipids, and an increased propensity for liver diseases (Voutilainen and Kärkkäinen 2019; Tilg, Cani, and Mayer 2016; Leclercq et al. 2024). People with AUD were distinguished from social drinkers and nondrinkers using metabolomics and identified differences in propionic acid and acetic acid in plasma (Mostafa et al. 2017). To date, there are few systematic analyses of sex-based differences in metabolomic profiles in humans because of low-powered samples or the lack of systematic data analysis (Costanzo et al. 2022).

Using preclinical models to determine the consequences of alcohol exposure during adolescence is a powerful approach due to ethical limiting factors of observational/survey-based research typically conducted in human studies. Therefore, preclinical models are essential as a tool and can serve as proxies for our understanding of alcohol-induced changes in human metabolomics (Humer, Probst, and Pieh 2020; Humer, Pieh, and Probst 2020). Plasma, urine, and fecal samples are minimally invasive sampling methods to determine alcohol-induced changes in metabolomic profiles (Mostafa et al. 2017; X. Wang et al. 2023). Previous research has used mouse models to determine early biomarkers of alcohol-induced liver disease (Manna et al. 2010; Manna, Thompson, and Gonzalez 2015; Manna et al. 2011; Bradford et al. 2008; Charkoftaki et al. 2022). Recently a rat model of AUD was established where relationships between changes in metabolome profiles and alcohol drinking and anxiety-like behaviors in adult male rats (X. Wang et al. 2023). However, to date there is limited work using mouse models to determine the link between affective behaviors and changes in metabolomic profiles. Therefore, in the present work, we determined short-term and long-term changes in metabolomic profiles and affective behaviors in male and female C57BL/6J mice following AIE exposure. Based on our previous work we expected greater changes in anxiety-like behavior in the female mice shortly after exposure and greater changes in anxiety-like behavior in male mice after long-term withdrawal. We also expected differences in metabolomic profiles of male and female AIE-exposed mice compared to air-exposed controls.

## 2. Methods

### 2.1 Subjects

Adolescent male and female C57BL/6J mice (n=10-13/group) were obtained from Jackson Laboratories (Bar Harbor, ME) on PND 21. All mice were allowed to acclimate to the colony for one week (PND 21-27) prior to experimentation, where they were handled daily for acclimation to experimenter manipulation. Mice were PND 28 at the beginning of the experiment. Animals were group-housed (4-5 per cage) with free access to food and water throughout the AIE or AIR exposure (detailed below). An experimental timeline is shown in **Figure 1**. All mice were maintained in a temperature and humidity-controlled room with lights on from 0700-1900 hr. Body weights were recorded each time the mice were exposed to the ethanol vapor inhalation chambers and each time they were tested in one of the behavioral tests. Animal care followed National Institutes of Health Guidelines under North Carolina Agricultural and Technical State University Institutional Animal Care and Use Committee approved protocols (21-005).

**Figure 1.**
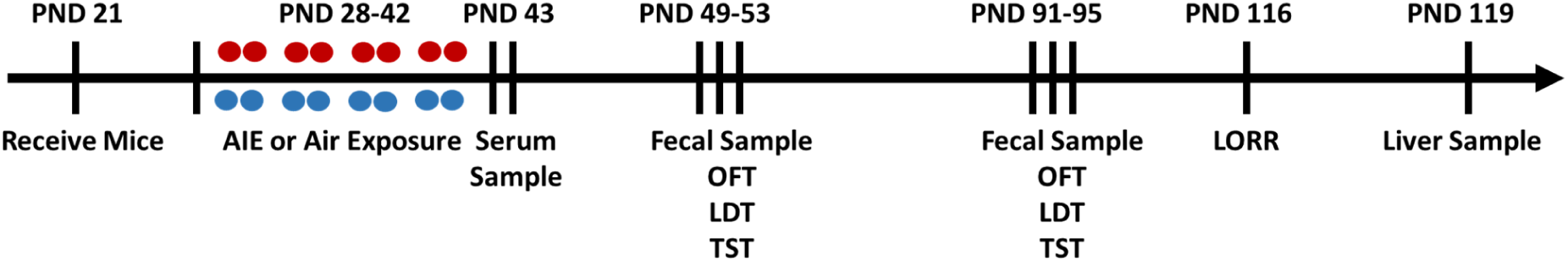
Experimental Timeline. Mice were exposed to AIE or AIR between PND 28-42. Red circles indicate ethanol exposure and blue circles indicate air exposure. A serum sample was collected on PND 42. Mice were run on consecutive days to the open field test (OFT), light/dark test (L/DT), and tail suspension test (TST) between PND 49-53 and again between PND 91-95. Fecal samples were collected during the OFT. Mice were challenged with 2.0 g/kg and tested for loss of righting reflex (LORR) on PND 116. Livers were extracted on PND 119.

### 2.2 Adolescent Intermittent Vapor Inhalation Chamber Exposure

Mice were exposed to repeated intermittent air or ethanol vapor for four two-day exposure cycles from PND 28-42 (Maldonado-Devincci et al. 2022). On PND 28-29, 32-33, 36-37, 40-41, at approximately 1630 hr, mice were weighed and administered an intraperitoneal injection (0.02 ml/g) of pyrazole (1 mmol/kg), an alcohol dehydrogenase inhibitor used to stabilize blood ethanol concentrations, combined with saline for control mice or combined with 1.6 g/kg ethanol (8% w/v) for ethanol-exposed mice and immediately placed in the inhalation chambers (23 in x 23 in x 13 in; Plas Labs, Lansing, MI). Mice remained in the vapor inhalation chambers for 16 hr overnight with room (control group) or ethanol (95% ethanol volatilized by passing air through an air stone (gas diffuser) submerged in ethanol) delivered to the chambers at a rate of 10 liters/min. The following morning at 0900 hr, mice were removed from the vapor inhalation chamber and 25μL of blood was collected from the submandibular space for blood ethanol concentration (BEC) assessment and then returned to the home cage for 8 hr. On PND 29, 33, 37, and 41, mice were administered pyrazole and placed in the chamber overnight for 16 hours. On intervening days, mice remained undisturbed in the home cage. All blood samples were centrifuged at 5,000*g* and serum was collected and used to analyze blood ethanol concentrations (Table 1) using the AM1 blood alcohol analyzer (Analox Instruments, Lunenburg, MA).

**Table 1:**
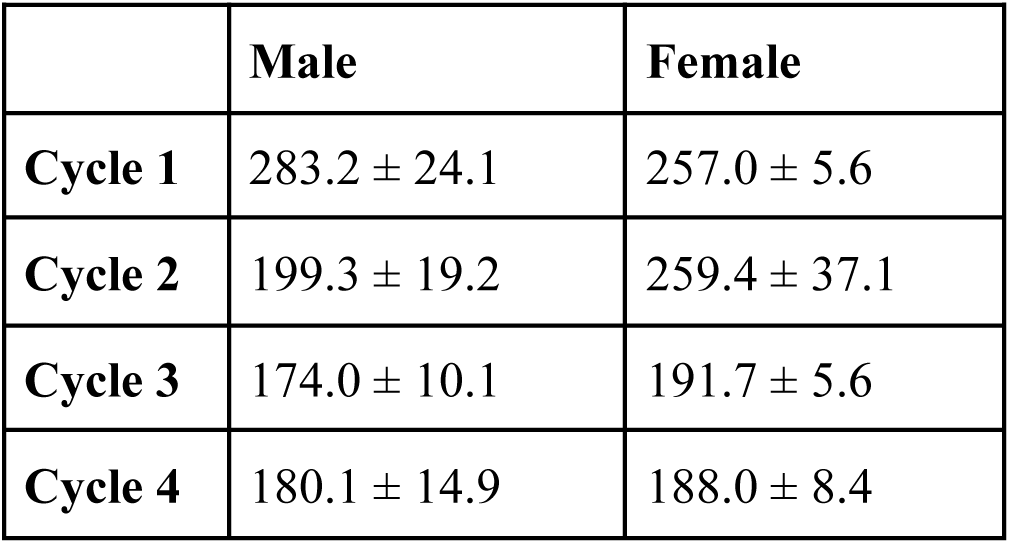
Blood ethanol concentrations (mg/dl) per cycle in male and female mice exposed to intermittent ethanol vapor inhalation. Data are presented as mean +/- SEM.

### 2.3 Behavioral Testing

Mice were tested over three consecutive days for behavioral changes using the open field test (OFT), light/dark test (LDT), and tail suspension test (TST) between PND 49-53 and again between PND 91-95. For all tests mice were counterbalanced across days and across groups for testing to minimize between-group differences in testing order. All mice were transported and acclimated to the behavioral testing room for at least 60 min before behavioral testing. All tests were conducted with experimenters blind to the sex and exposure conditions of the mice. Experimental details are described below.

#### 2.3.1 Open Field Test

Between PND 49-51 and again on PND 91-93, mice were tested for a 60-minute trial in the open field test (Tatem et al. 2014). Mice were tested with regular overhead lighting (400 lux). Using a Plexiglas chamber (40.6 cm x 40.6 cm), mice were introduced facing one of the corners selected at random to the open field testing arena (Kinder Scientific, Poway, CA) to assess general exploratory behavior (distance traveled and rearing) and activity in the center zone as a measure of anxiety-like behavior (latency, time spent, and distance traveled in the center zone). Immediately upon removal from the arena, mice were returned to their home cage and returned to the colony. The open field arena was cleaned with 70% ethanol and allowed to dry completely before the next mouse was introduced. Data were captured through beam breaks using the SmartFrame Open Field System Data and were quantified using Motor Monitoring Behavioral software (Kinder Scientific, Poway, CA).

#### 2.3.2 Light/Dark Test

The Light/Dark Test (LDT) is used to evaluate anxiolytic behavior in rodents as they have a natural aversion to brightly lit environments (Takao and Miyakawa 2006). With this test, anxiety in mice can be accurately expressed through their locomotor activity and time spent in each side of the apparatus (Himanshu et al. 2020). On PND 50-52 and 93-94 the LDT was conducted for 10 min using the same chamber as the open field test with an infrared transparent black divider (20.3 cm x 20.3 com) inserted into the dark compartment (Kinder Scientific, Poway, CA). The light side of the box was illuminated with a bright light (850 lux). Mice were introduced to the dark compartment and latency to enter and time spent on the light side were quantified along with distance traveled in both compartments. After each trial, the chamber was sanitized with 70% alcohol and allowed to completely dry. Data were captured through beam breaks using the SmartFrame Open Field System and data were quantified using Motor Monitoring Behavioral software (Kinder Scientific, Poway, CA).

#### 2.3.3 Tail Suspension Test

The tail suspension test (TST) attempts to translate depressive-like behaviors in humans by observing the initial escape oriental movements made when the rodent is placed in an inescapable environment (Cryan, Mombereau, and Vassout 2005). The TST assessment was conducted between PND 51-53 and again between PND 93-95 to test the longitudinal effects of ethanol exposure and withdrawal on depressive-like behaviors. The TST was conducted using previously established procedures (Can et al., 2012). After acclimation mice were briefly restrained to pass the tail through a small plastic cylinder to prevent tail climbing and then attach the adhesive strip to the end of the tail. Mice were then suspended 60 cm above the ground for a duration of 6 min. Mice were then immediately returned to their home cage after the adhesive strip and plastic cylinder were removed. Sessions were recorded and scored offline by experimenters’ blind to exposure and sex conditions for latency to immobility and duration of immobility. Data were quantified for time spent immobile using previously established protocols described by (Can et al. 2012).

### 2.4 Loss of Righting Reflex

On PND 116, all mice were challenged with 4.0 g/kg ethanol and assessed for loss of righting reflex (LORR) according to previously published methods (Silvers et al. 2003; Maldonado-Devincci and Kirstein 2020). Briefly, mice were administered an i.p. injection of ethanol (20% v/v) and placed in a supine position in a polystyrene reagent reservoir. The duration of time it took (duration of LORR) for the mouse to right itself on all four paws for at least 30 seconds was recorded. Any mouse that did not lose its righting reflex was not included in the analysis (Male-Air n=1 of 10, Male-Ethanol n=7 of 10), Female-Air n=2 of 10, Female-Ethanol n=5 of 10). Upon recovery of LORR, blood was collected from the submandibular space and used to measure blood ethanol concentrations. After blood collection, mice were returned to their home cage and allowed to fully recover from intoxication. Mice were left in the home cage until they were euthanized on PND 119 when liver samples were collected for metabolite analysis. All blood samples were centrifuged at 5000 g and serum was collected to analyze blood ethanol concentrations using the AM1 Analox Alcohol Analyzer (Analox Instruments, Lunenburg, MA, USA). During processing some of the samples were lost due to technical difficulties.

### 2.5 Metabolomics

A metabolomic nuclear magnetic resonance (NMR) assay was performed on serum samples collected on PND 43 and fecal samples from PND 49-51 and PND 91-93 following the open field test. All samples (serum, fecal, and tissue) were frozen at -80°C until processed for analysis. Samples were thawed on ice before the NMR analysis. Fecal samples were extracted using water based on a previously reported method with slight changes (Gratton et al. 2016). These extract samples were mixed with a phosphate buffer of D2O which made the the final samples contain 10% of D2O with 0.1 M phosphate buffer (pH = 7.4) and 0.5 mM trimethylsilyl propanoic acid (TSP). The samples were transferred to 5 mm NMR tubes after being centrifuged for further nuclear magnetic resonance (NMR) acquisition.

The serum samples were then extracted using a NaCl approach followed by a previously reported approach. The liver tissue was extracted using a two-step method (Wu et al. 2008) including the homogenization of tissue in cold 2.5:1 methanol-water solvent followed by addition of ice-cold chloroform and water solvent. After centrifugation, the upper polar phase was dried and reconstituted in a phosphate buffer. The final sample phosphate and TSP concentrations were the same as the fecal samples.

### 2.6 Design and Analyses

Behavioral data were analyzed using a three-factor mixed-model design ANOVA with Exposure (2; Air, Ethanol) and Sex (2; Male, Female) as between-subjects factors and Age as a repeated measure. Blood ethanol concentrations during AIR or AIR exposure were analyzed using a two-factor mixed model ANOVA with Sex (2; Male, Female) as a between subject’s factor and Cycle as a repeated measure. LORR data were analyzed using a two-factor between-subjects design ANOVA for Sex and Exposure. Apriori comparisons of exposure were also conducted within each sex for each behavioral measure. In the presence of significant interactions, Tukey’s and Sidak’s multiple comparisons post hoc tests were used where appropriate. Data were analyzed with GraphPad Prism version 10.4.1 for Windows, GraphPad Software, Boston, Massachusetts USA.

The NMR spectra were preprocessed using Bruker Topspin 4.11 and carefully checked in Bruker Amix 4.0 before peak bucketing. The NMR peaks bucketing were conducted using previously established protocols (Wang, Maldonado-Devincci, and Jiang 2020) with slight changes. The processed data were normalized to the total peak intensity exported to Excel (Microsoft). Metabolite identification was carried out using Chenomx 8.4 (Chenomx Inc). The Student two-way ANOVA (two tails) were calculated in Excel (Microsoft). The principal component analysis (PCA) was carried out using PLS-toolbox (Eigenvector Research). The box plots and heatmap were plotted using MetaboAnalyst 5.0.

## 3. Results

### 3.1 Blood ethanol concentrations

Blood ethanol concentrations (Table 1) varied across cycles (F_(3, 53)_=13.83, p<0.0005). However, post hoc tests failed to reveal any statistically significant differences between cycles. There were no differences in BECs between males and females at any given exposure cycle (F_(1, 18)_=0.7304, p>0.05).

### 3.2 Behavioral Assessment

Mice were tested for behavioral changes at two time points, shortly after AIE exposure (PND 49-53) and again in adulthood (PND 91-95). All mice underwent a series of behavioral tests over three consecutive days. This included the open field (**Table 2**), light/dark (**Table 3**), and tail suspension **(Table 4**) tests to determine changes in exploratory behavior, anxiety-like behavior, and depressive-like behavior, respectively.

**Table 2:** Open Field Test Statistics.

|  | Open Field Test |  | Rearing |  |
| --- | --- | --- | --- | --- |
| Factor | F | p | F | p |
| Trial | $F(1,43) = 3.92$ | 0.054 | $F(1,43) = 0.01$ | 0.943 |
| Sex | $F(1,43) = 0.27$ | 0.603 | $F(1,43) = 1.64$ | 0.207 |
| Exposure | $F(1,43) = 0.03$ | 0.849 | $F(1,43) = 3.93$ | 0.054 |
| Trial x Sex | $F(1,43) = 0.85$ | 0.361 | $F(1,43) = 2.37$ | 0.131 |
| Trial x Exposure | $F(1,43) = 0.15$ | 0.696 | $F(1,43) = 4.72$ | 0.035 |
| Sex x Exposure | $F(1,43) = 2.06$ | 0.159 | $F(1,43) = 0.78$ | 0.381 |
| Trial x Sex x Exposure | $F(1,43) = 1.26$ | 0.268 | $F(1,43) = 0.99$ | 0.324 |

**Table 3:**
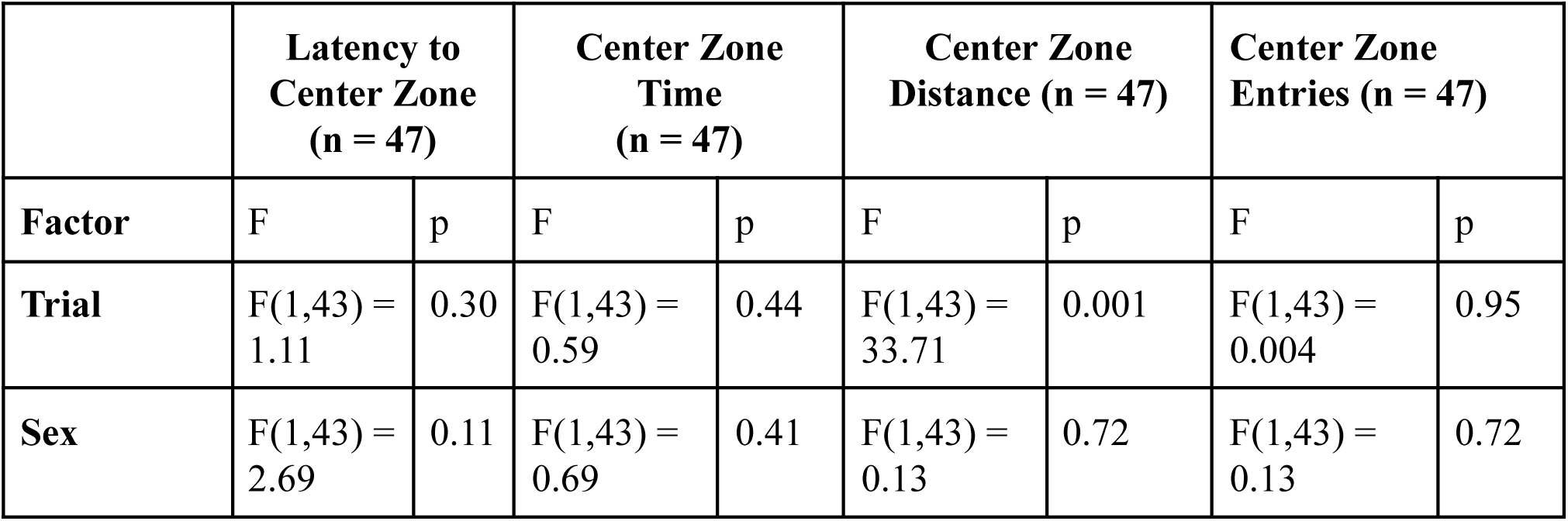
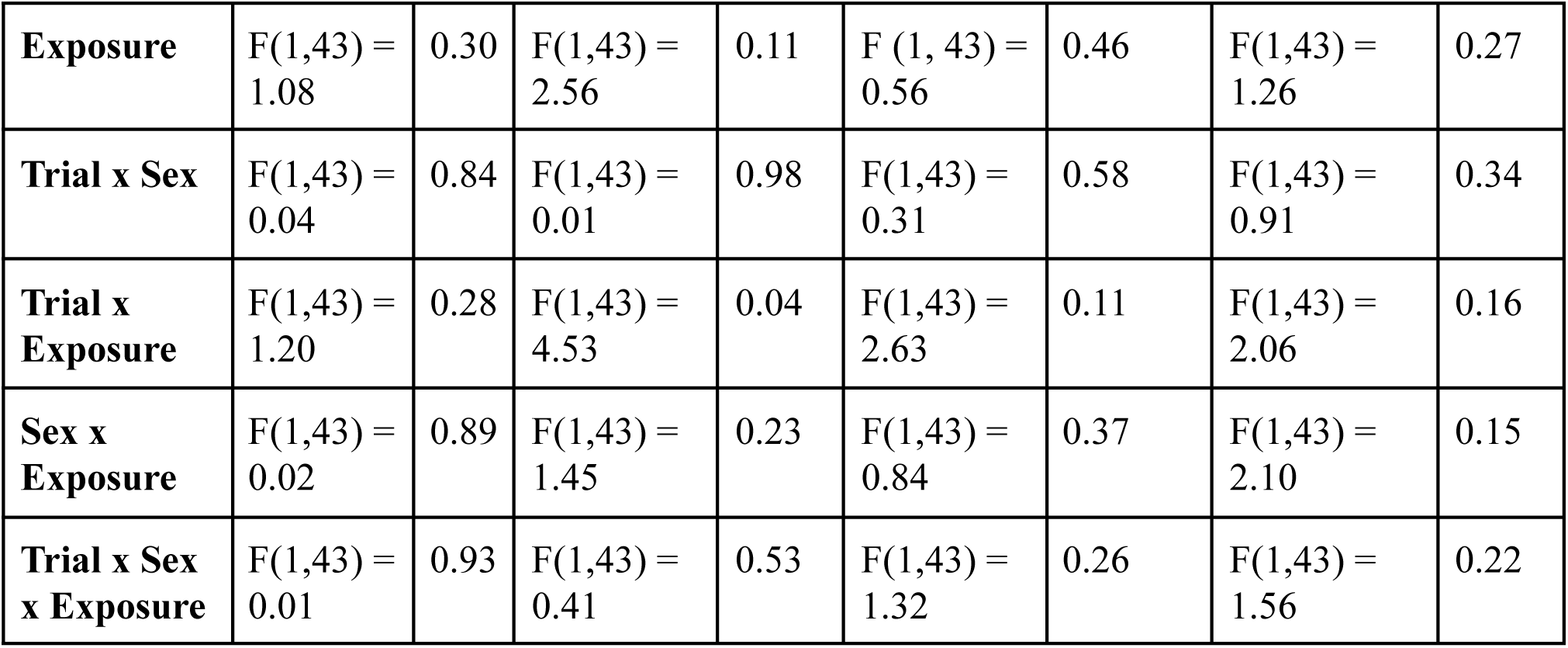
Center Zone Statistics.

**Table 4:** Light Dark Test Statistics.

|  | **Latency to Enter Light** |  | **Light Zone Distance (% of total)** |  | **Light Side Entries** |  | **Time in Light Side (%)** |  |
| --- | --- | --- | --- | --- | --- | --- | --- | --- |
| **Factor** | F | p | F | p | F | p | F | p |
| **Trial** | F(1,43) = 0.23 | 0.64 | F(1,43) = 8.88 | 0.005 | F(1,43) = 35.68 | 0.001 | F(1,43) = 17.22 | <0.001 |
| **Sex** | F(1,43) = 9.8 | 0.003 | F(1,43) = 0.58 | 0.45 | F(1,43) = 0.99 | 0.32 | F(1,43) = 0.56 | 0.46 |
| **Exposure** | F(1,43) = 0.45 | 0.51 | F(1,43) = 0.33 | 0.57 | F(1,43) = 0.28 | 0.60 | F(1,43) = 0.79 | 0.38 |
| **Trial x Sex** | F(1,43) = 0.41 | 0.52 | F(1,43) = 1.55 | 0.22 | F(1,43) = 0.04 | 0.84 | F(1,43) = 0.68 | 0.41 |
| **Trial x Exposure** | F(1,43) = 0.02 | 0.88 | F(1,43) = 0.63 | 0.43 | F(1,43) = 0.01 | 0.94 | F(1,43) = 0.68 | 0.42 |
| **Sex x Exposure** | F(1,43) = 0.14 | 0.71 | F(1,43) = 0.28 | 0.60 | F(1,43) = 0.81 | 0.37 | F(1,43) = 0.74 | 0.40 |
| **Trial x Sex x Exposure** | F(1,43) = 0.08 | 0.79 | F(1,43) = 1.55 | 0.22 | F(1,43) = 1.32 | 0.26 | F(1,43) = 0.31 | 0.59 |

#### 3.2.1 Open Field Test

Overall, there were no changes in distance traveled following AIE and short-term withdrawal (PND 49) or long-term withdrawal (PND 91) in male or female mice (**Figure 2A**; **Table 2).** Male AIE-exposed mice showed higher rearing at both time points compared to air-exposed mice. However, females did not show any differences in rearing behavior at either time point (**Figure 2B**). There were no differences in latency to enter the center zone, distance traveled in the center zone (**Figure 2D)**, or entries to the center zone (**Table 3**). However, male ethanol-exposed mice spent more time in the center zone compared to air-exposed controls; which was primarily driven by the later time point. However, there were no differences observed in females at either time point (**Figure 2C**; **Table 3**).

**Figure 2.**
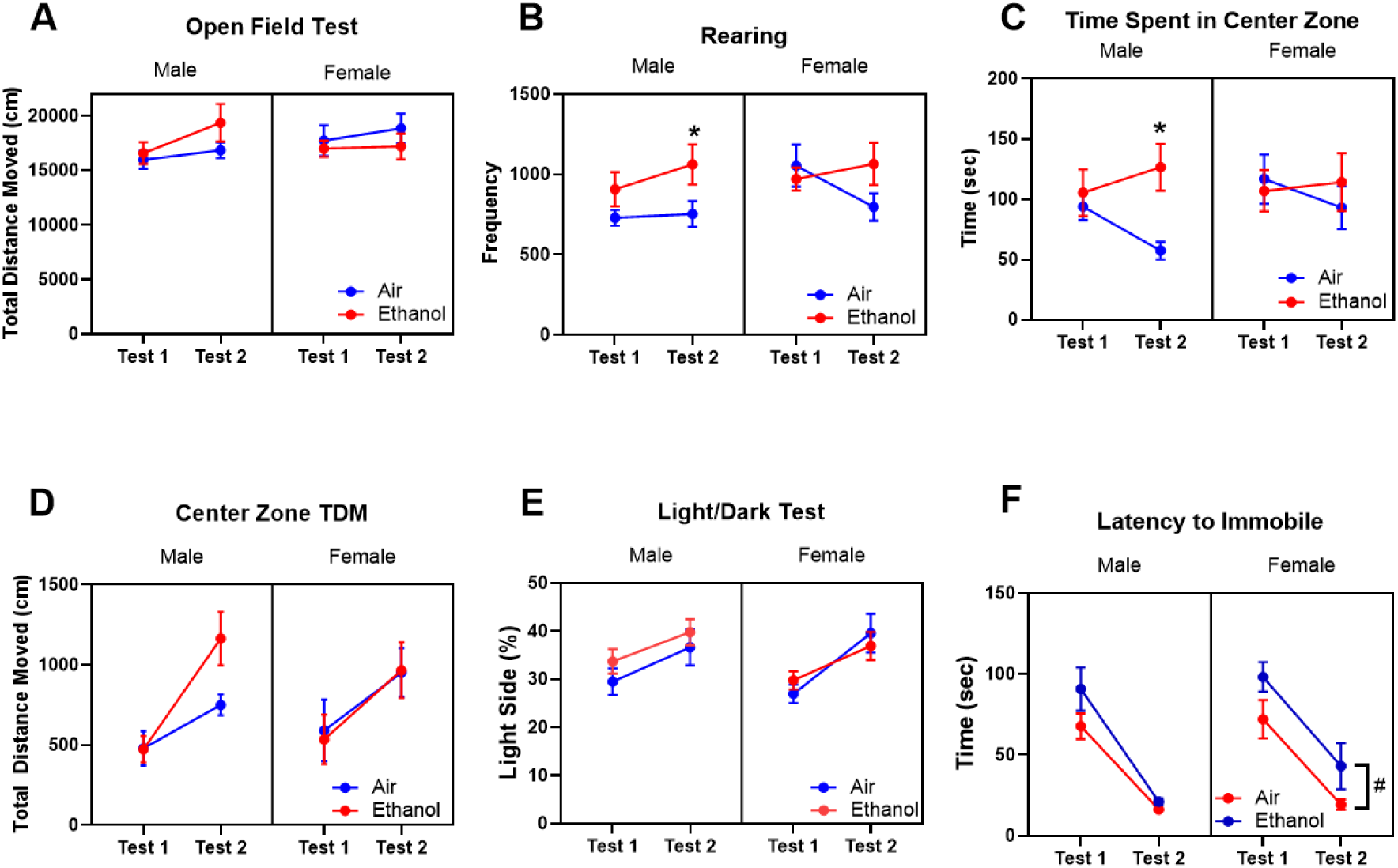
Behavioral Measures. Overall behavior was mildly altered after AIE exposure and short-term (PND 49-51) and long-term (PND 91-93) withdrawal. (A) Total distance traveled in the entire open field arena; (B) Rearing in the open field test; (C) Time spent in the center zone in the open field test; (D) Distance Traveled in the open field test; (E) percent time spent on the light side of the light dark test; (F) latency to become immobile in the tail suspension test. Data are presented as mean +/- SEM. * indicates a significant difference between ethanol-exposed and air-exposed mice. # indicates a main effect of exposure.

For the light/dark test (LDT), male mice took longer to enter the light compartment compared to females, regardless of pre-exposure. There were no differences between groups in distance traveled in the light compartment, number of entries into the light compartment, or percent time spent in the light compartment (**Figure 2E**; **Table 4**).

For the tail suspension test, mice became immobile faster (**Figure 2F)** on the second trial compared to the first trial. For female mice, AIE-exposed mice took longer to become immobile compared to AIR-exposed mice at both time points (main effect of exposure). However, this effect was absent in males. There were no differences in the time spent immobile at either time point, however, both males and females spent more time immobile at the later time point compared to the earlier time point (**Table 5**).

**Table 5:** Tail Suspension Test Statistics.

| **Factor** | **Latency to Immobile** |  | **Duration Immobile** |  |
| --- | --- | --- | --- | --- |
|  | F | p | F | p |
| **Trial** | F (1, 43) = 119.4 | <0.001 | F(1,43) = 207.9 | <0.001 |
| **Sex** | F (1, 43) = 1.44 | 0.24 | F(1,43) = 1.22 | p = 0.28 |
| **Exposure** | F (1, 43) = 6.44 | 0.01 | F(1,43) = 0.86 | p = 0.36 |
| **Trial x Sex** | F (1, 43) = 0.40 | 0.53 | F(1,43) = 0.29 | p = 0.59 |
| **Trial x Exposure** | F (1, 43) = 0.95 | 0.33 | F(1,43) = 0.12 | p = 0.73 |
| **Sex x Exposure** | F (1, 43) = 0.52 | 0.48 | F(1,43) = 0.05 | p = 0.83 |
| **Trial x Sex x Exposure** | F (1, 43) = 0.57 | 0.46 | F(1,43) = 0.52 | p = 0.47 |

### 3.4 Metabolomics-Metabolite analyses

Supplemental results showing correlations between behavior and specific metabolites are shown in the supplemental section.

#### 3.4.1 Serum Metabolites

The serum samples were collected 24 hours after the last exposure cycle on PND 43 for all mice. The PCA score plot showed that the male samples (**Figure 3A**) had better separation than the female samples (**Figure 3B**). These results indicate that the metabolic profiling was highly impacted by ethanol exposure in the male samples, which we interpret as these samples being more distinct than the female samples. The PCA loadings showed that amino acids and isoleucine [Exposure (F_1, 20_ = 11.29, p = 0.003); **Figure 4C**], leucine [Exposure (F_1, 20_ = 27.10, p < 0.0001); **Figure 4D**] , and valine [Exposure (F_1, 20_ = 30.67, p < 0.0001); **Figure 4J**] contributed highly to the separation of air-control samples from the ethanol-exposed samples, all of which were significantly downregulated after ethanol exposure. The results also indicate the potential high demand of those amino acids in the blood. Asparagine [Sex by Exposure (F_1, 20_ = 4.53; p < 0.05); **Figure 4A**], glucose [Exposure (F_1, 20_ = 7.71, p < 0.0030); **Figure 4B**], lipid [Exposure (F_1, 20_ = 8.54, p < 0.01); Sex (F_1, 20_ = 13.73, p < 0.002); **Figure 4E**] and lipoprotein [Sex (F_1, 20_ = 9.94, p < 0.006); Exposure (F_1, 20_ = 3.91, p = 0.06); Sex by Exposure (F_1, 20_ = 3.46, p = 0.07); **Figure 4F**] were also affected by ethanol exposure in the male group, but the changes in the female group were much weaker.

**Figure 3.**
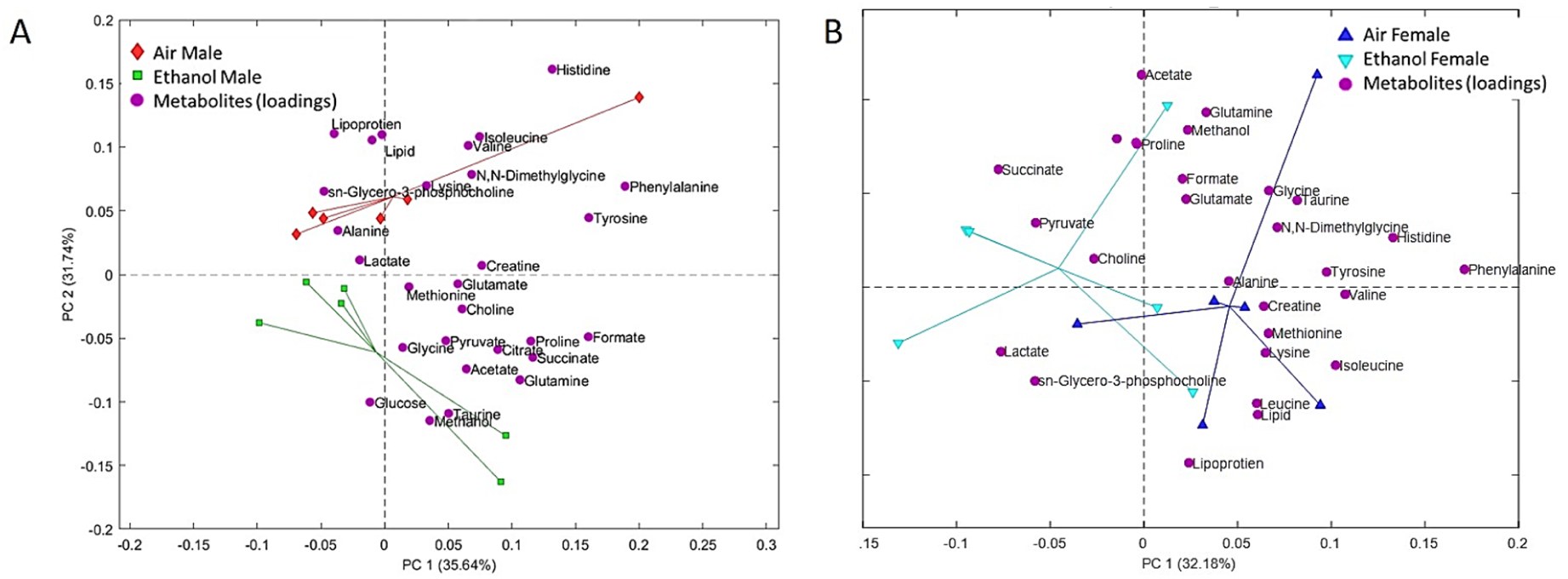
PCA of serum samples. (A) PCA results from male serum samples. Red diamonds are Air-Male, Green squares are Ethanol-Male mice. (B) PCA results from female serum samples. Blue upward triangles are Air-Female, and Cyan downward triangles are Ethanol-Female mice. Magenta circles (both panels) are the loading samples which are the contributions of metabolites with the PCA model.

**Figure 4.**
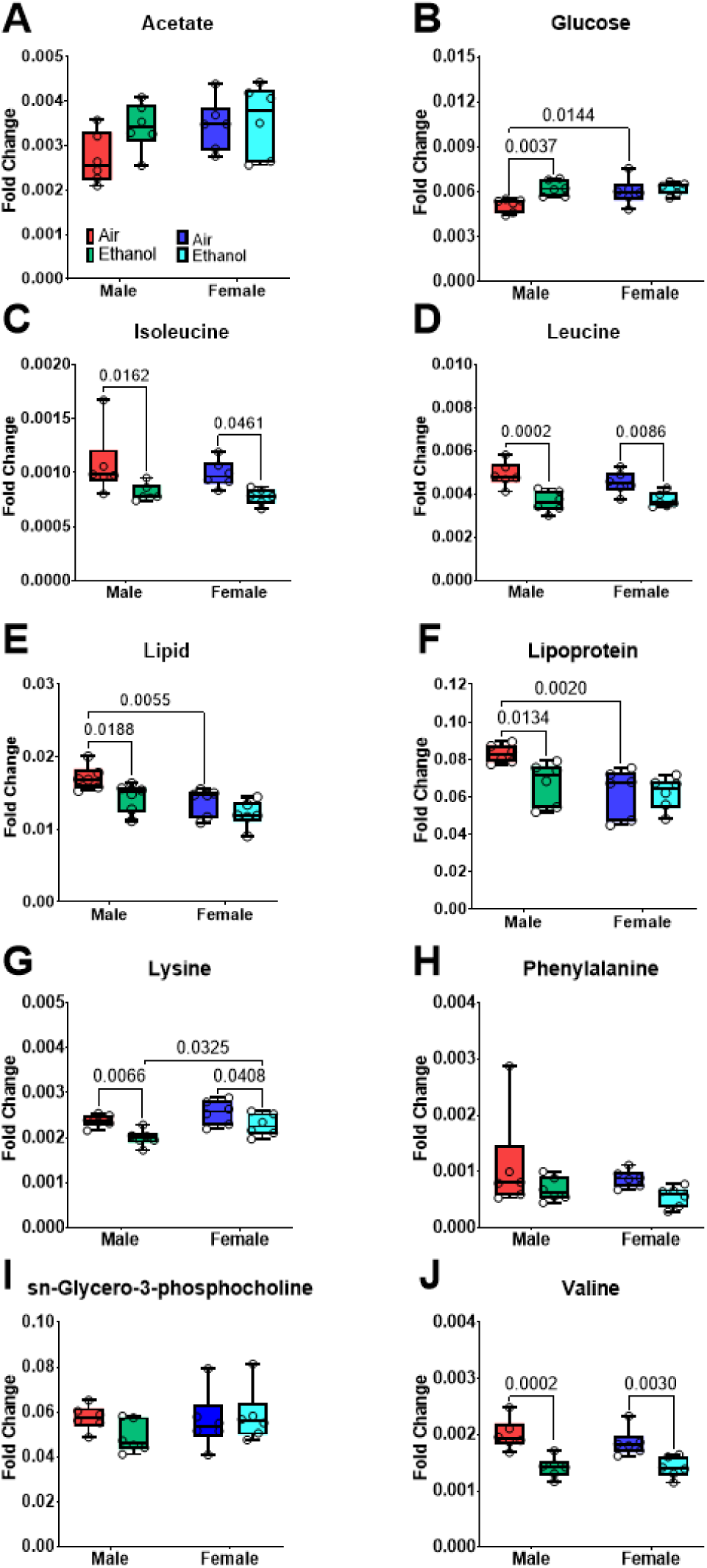
Selected metabolic changes in both male and female serum samples collected on PND 43. The samples include male control (Red), male ethanol (Green), female control (Blue) and female ethanol (Cyan).Data are represented as box and whisker plots for median and range. All data points are shown as open circles.

#### 3.4.2 Fecal Metabolites-short term withdrawal

Short-term withdrawal fecal samples were collected between PND 49-51, which was approximately 1 week after the last cycle of AIE exposure. The metabolic profiling showed distinct differences between the air- and ethanol-exposed male mice (**Figure 5A**). However, the female air- and ethanol-exposed mice only showed subtle differences in metabolic profiles after AIE (**Figure 5B**). The primary metabolites that contributed to the separation in male ethanol- and air-exposed mice include amino acids such as isoleucine [Exposure (F_1, 20_ = 11.36, p < 0.004); Sex (F_1, 20_ = 13.54, p < 0.002); **Figure 6F**] and leucine [Sex by Exposure (F_1, 20_ = 6.26, p < 0.03); Exposure (F_1, 20_ = 12.45, p < 0.003); **Figure 6G**]. Males showed ethanol-induced separation in metabolite profiles for amino acids including asparagine [Sex by Exposure (F_1, 20_ = 4.53, p < 0.05); **Figure 6B**] and valine [Exposure (F_1, 20_ = 5.18, p < 0.04); **Figure 6J**]. Other metabolites including cholate [Sex by Exposure (F_1, 20_ = 10.06; p < 0.005); Sex (F_1, 20_ = 10.06, p < 0.005); Exposure (F_1, 20_ = 32.27, p < 0.001); **Figure 6C**], ethanolamine [Sex by Exposure (F_1, 20_ = 6.56, p < 0.02); Exposure (F_1, 20_ = 12.50, p < 0.003); **Figure 6D**], glycocholate [Sex by Exposure (F_1, 20_ = 7.68, p < 0.02); Sex (F_1, 20_ = 8.37, p < 0.01); Exposure (F_1, 20_ = 6.15, p < 0.03); **Figure 6E**], and saccharopine [Sex by Exposure (F_1, 20_ = 8.96, p < 0.01); Sex (F (1, 20) = 17.97, p < 0.0005); Exposure (F_1, 20_ = 4.77, p < 0.05); **Figure 6H**] are also important for the separations in overall profiles for the male and female mice.

**Figure 5.**
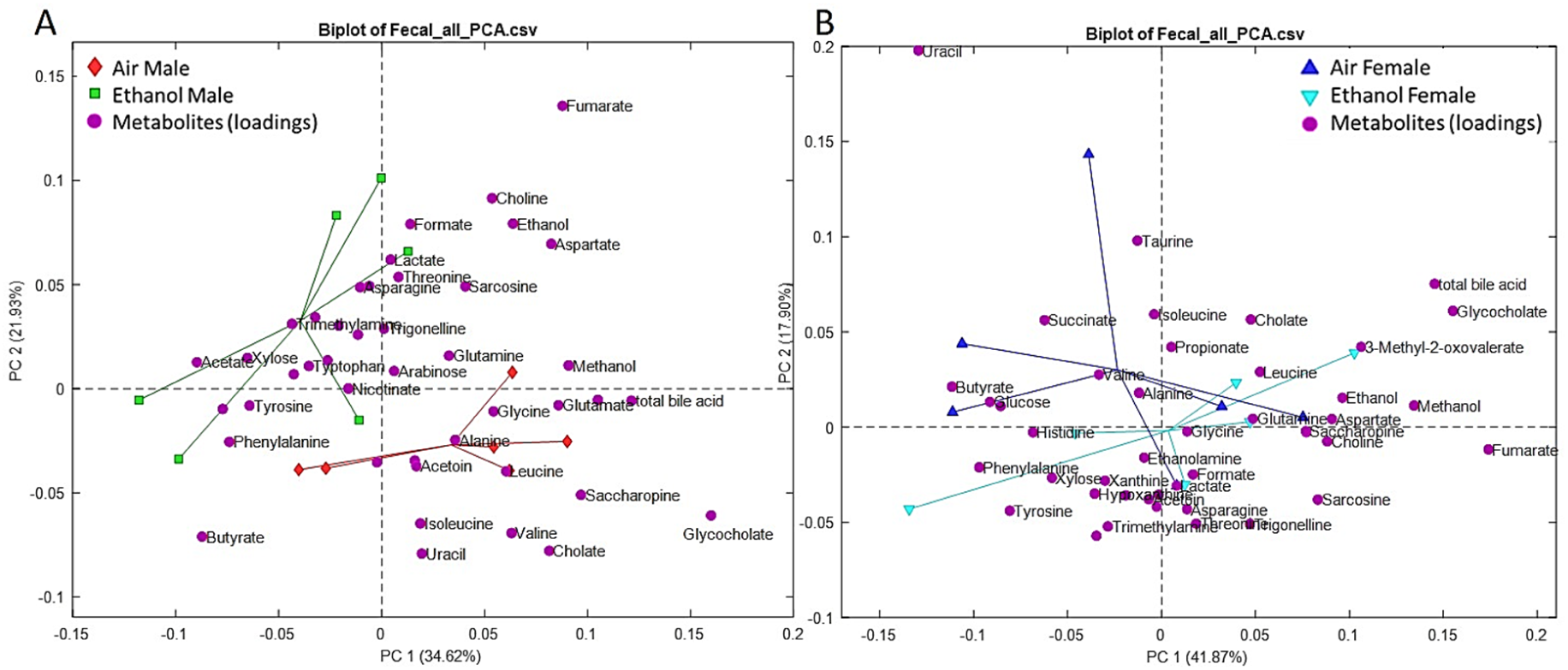
The PCA studies of short-term withdrawal fecal metabolites. (A) PCA results of male fecal samples. Red diamonds are Air-Male, Green squares are Ethanol-Male. (B) PCA results for female fecal samples. Blue upward triangles are Air-Female, and Cyan downward triangles are Ethanol-Female mice. Magenta circles are the loading samples which are the contributions of metabolites with the PCA model (both panels).

**Figure 6:**
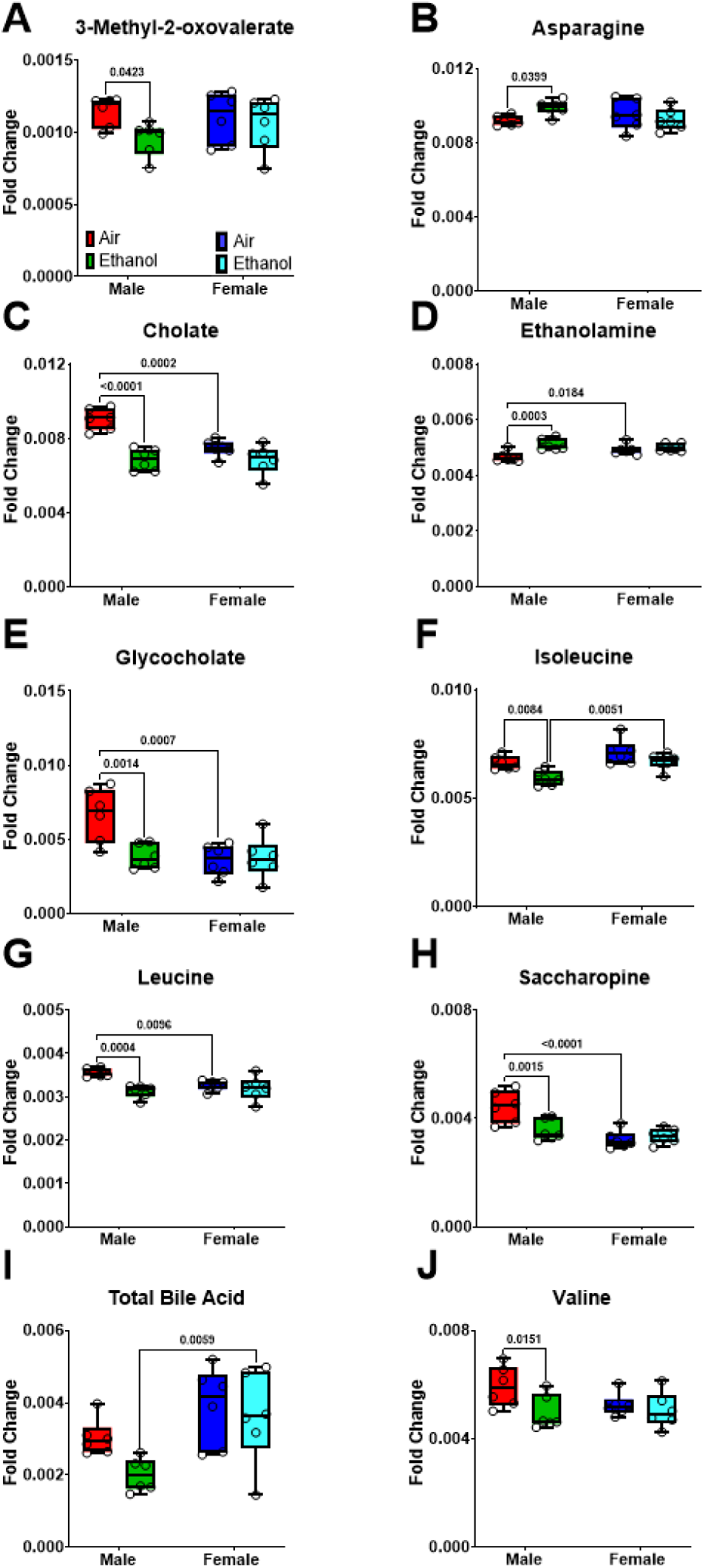
Selected metabolites for fecal short-term withdrawal following AIE. The samples include male control (Red), male ethanol (Green), female control (Blue) and female ethanol (Cyan). Data are represented as box and whisker plots for median and range. All data points are shown as open circles.

#### 3.4.3 Fecal Metabolites-Long-term withdrawal

In the long term withdrawal analyses with fecal samples that were collected between PND 91-95 (**Figure 7)**, males showed a clear partial separation between air- and ethanol-exposed groups (**Figure 7A)**; but the females showed mixed results with greater overlap in the metabolic profiles between air- and ethanol-exposed mice (**Figure 7B**). Sex differences were observed in overall metabolic profiles regardless of exposure for cholate [Sex (F_1, 18_ = 15.37, p < 0.0002); **Figure 8C**], glycocholate [Sex (F_1, 18_ = 25.10, p < 0.0001); **Figure 8E**], saccharopine [Sex (F_1, 18_ = 45.00, p < 0.0001), **Figure 8H**], total bile acid [Sex (F_1, 18_ = 5.99, p < 0.03), **Figure 8I**], and valine [Sex (F_1, 18_ = 9.69, p < 0.01), **Figure 8J**]. There were no ethanol-induced separations in overall fecal metabolite profiles in male or female mice following long-term withdrawal after AIE exposure.

**Figure 7.**
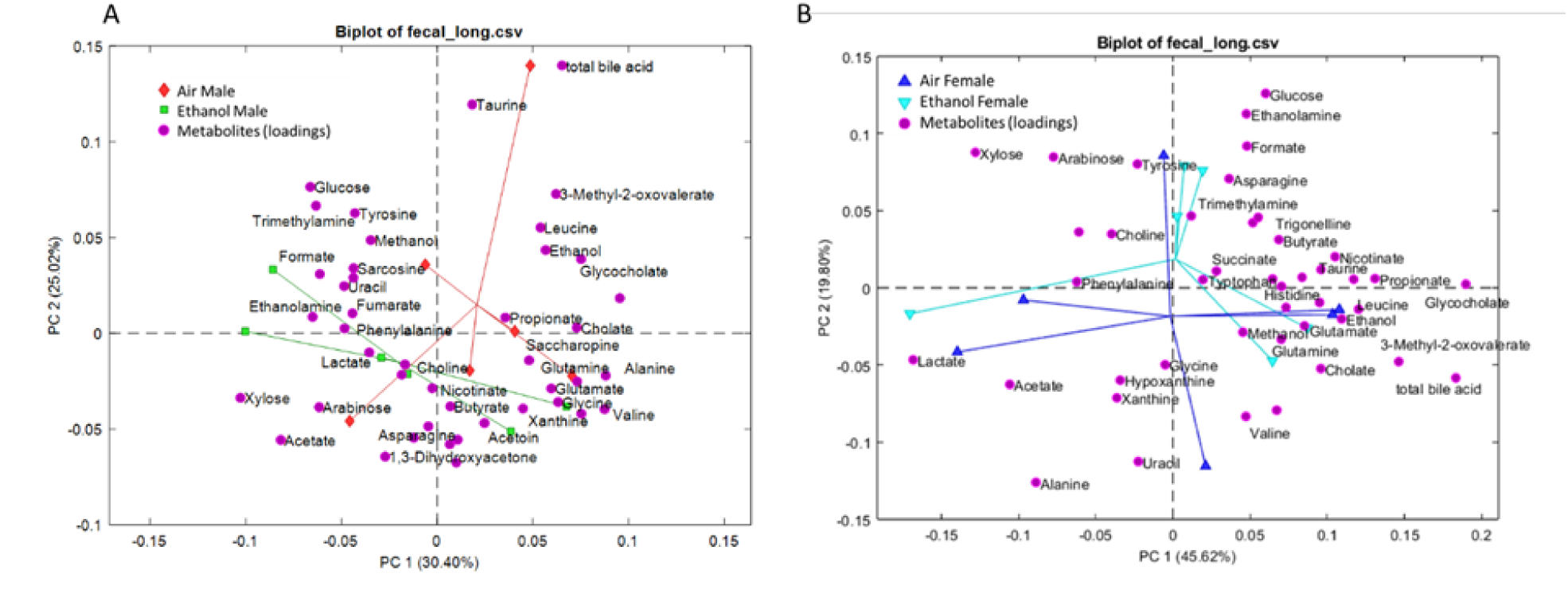
The PCA studies of long-term withdrawal fecal metabolites. (A) PCA results of male fecal samples. Red diamonds are Air-Male, Green squares are Ethanol-Male, (B) PCA results of female fecal samples. Blue upward triangles are Air-Female, and Cyan downward triangles are Ethanol-Female. Magenta circles represent the loading samples which are the contributions of metabolites with the PCA model (both panels).

**Figure 8.**
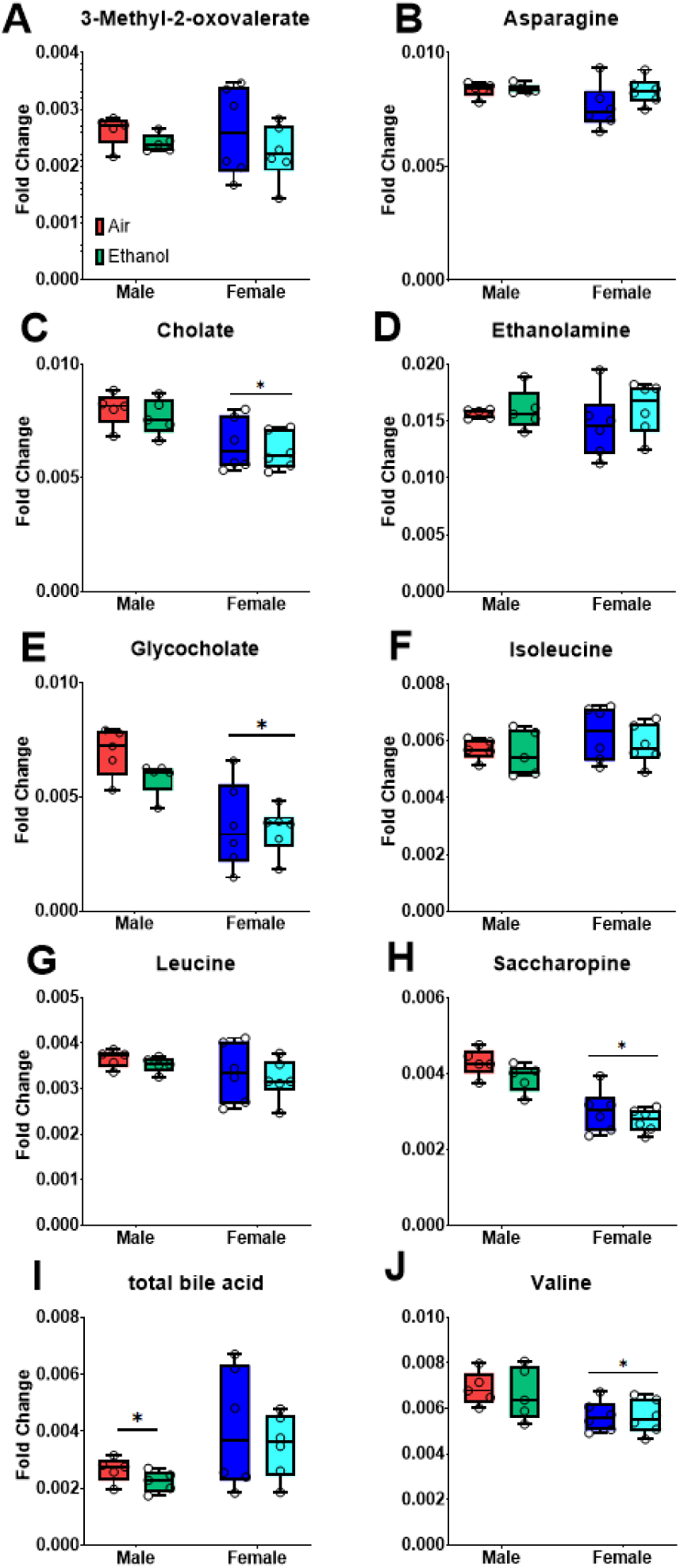
Selected metabolites for fecal long-term withdrawal samples following AIE. The samples include male control (Red), male ethanol (Green), female control (Blue) and female ethanol (Cyan). Data are represented as box and whisker plots for median and range. All data points are shown as open circles.

#### 3.4.4 Liver Metabolites

Three days following the LORR test (detailed below), liver samples were collected on PND 119. It is important to note that all mice were exposed to the sedating ethanol dose for the LORR test before liver samples were collected and analyzed for changes in metabolite profiles. The liver metabolites did not show clear separation in the PCA score plot in both male and female samples which indicated no high metabolic perturbation in the liver metabolites. Male and female are still different and the ethanol and air are closer to the female group, but the separation is just partial (**Figure 9**). Most of the differences in liver metabolite profiles showed sex differences between males and females, regardless of ethanol exposure including choline [Sex (F_1, 20_ = 7.81, p < 0.02); **Figure 10A**], glucose [Sex (F_1, 20_ = 5.16, p < 0.05); **Figure 10B**], glutamate [Sex (F_1, 20_ = 4.961, p < 0.05); **Figure 10C**], isocitrate [Sex (F_1, 20_ = 6.02, p < 0.03); **Figure 10F**], nicotinurate [Sex (F_1, 20_ = 6.02, p < 0.03); **Figure 10G**], phenylalanine [Sex (F_1, 20_ = 4.40, p < 0.05); **Figure 10H**], and UMP [Sex (F_1, 20_ = 4.69, p < 0.05); **Figure 10J**]. Female mice exposed to ethanol had higher levels of glycine compared to controls and their male-exposed counterparts [Sex by Exposure (F_1, 20_ = 6.06, p < 0.03); Sex (F_1, 20_ = 6.16, p < 0.03); **Figure 10D**]. In contrast, ethanol increased taurine levels only in ethanol-exposed male mice compared to controls [Sex by Exposure (F_1, 20_ = 4.56, p < 0.05); **Figure 10I**].

**Figure 9:**
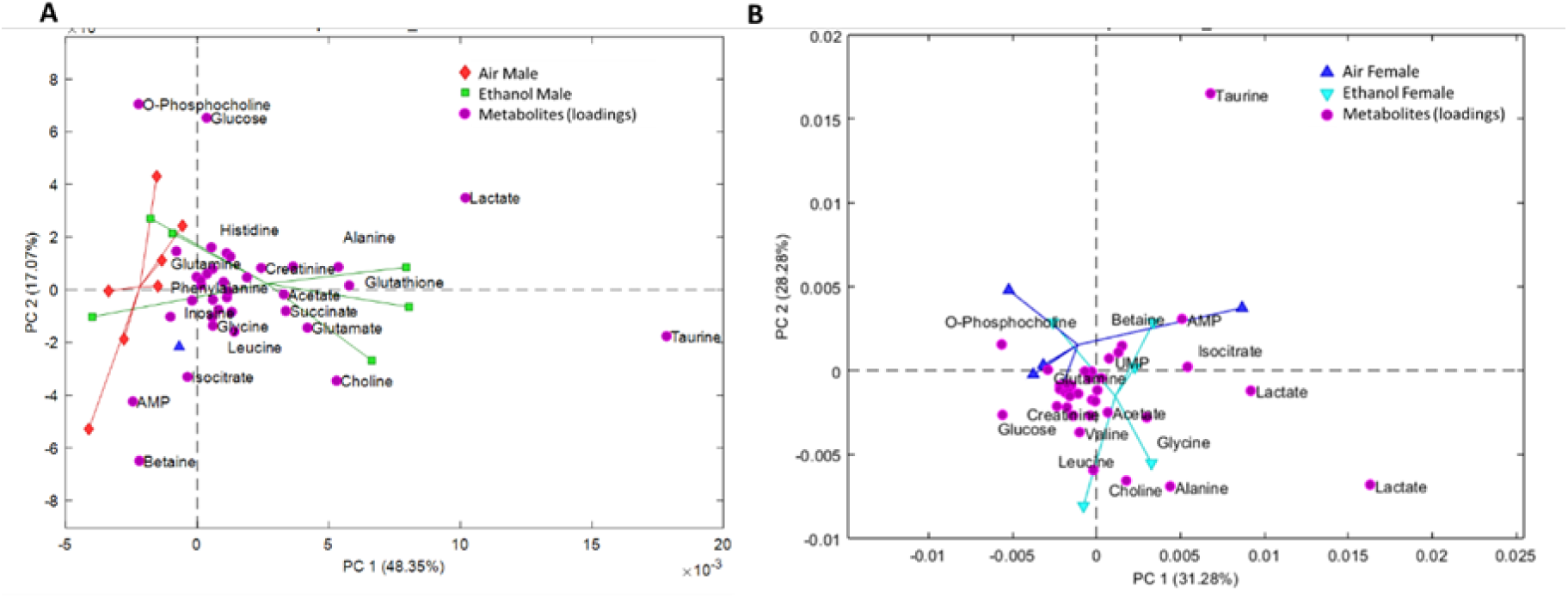
The PCA studies of liver metabolites. (A) PCA results for male liver samples. Red diamonds are Air-Male, Green squares are Ethanol-Male mice. (B) PCA results for female liver samples. Blue upward triangles are Air-Female, and Cyan downward triangles are Ethanol-Female. Magenta circles are the loading samples which are the contributions of metabolites with the PCA model (both panels).

**Figure 10.**
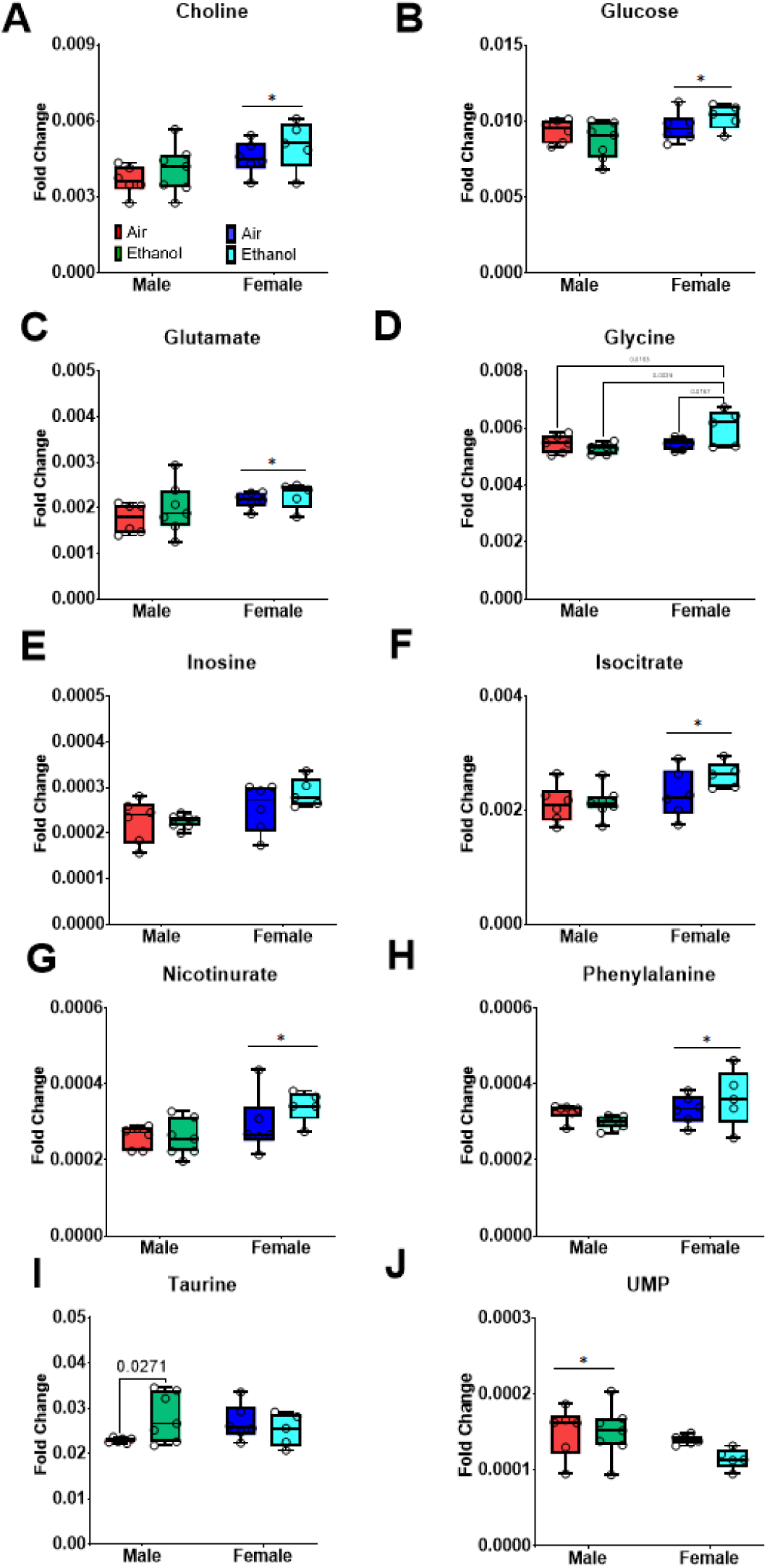
The PCA studies of liver metabolites: Selected metabolites for liver samples. The samples include male control (Red), male ethanol (Green), female control (Blue) and female ethanol (Cyan). Data are represented as box and whisker plots for median and range. All data points are shown as open circles.

**Figure 11.**
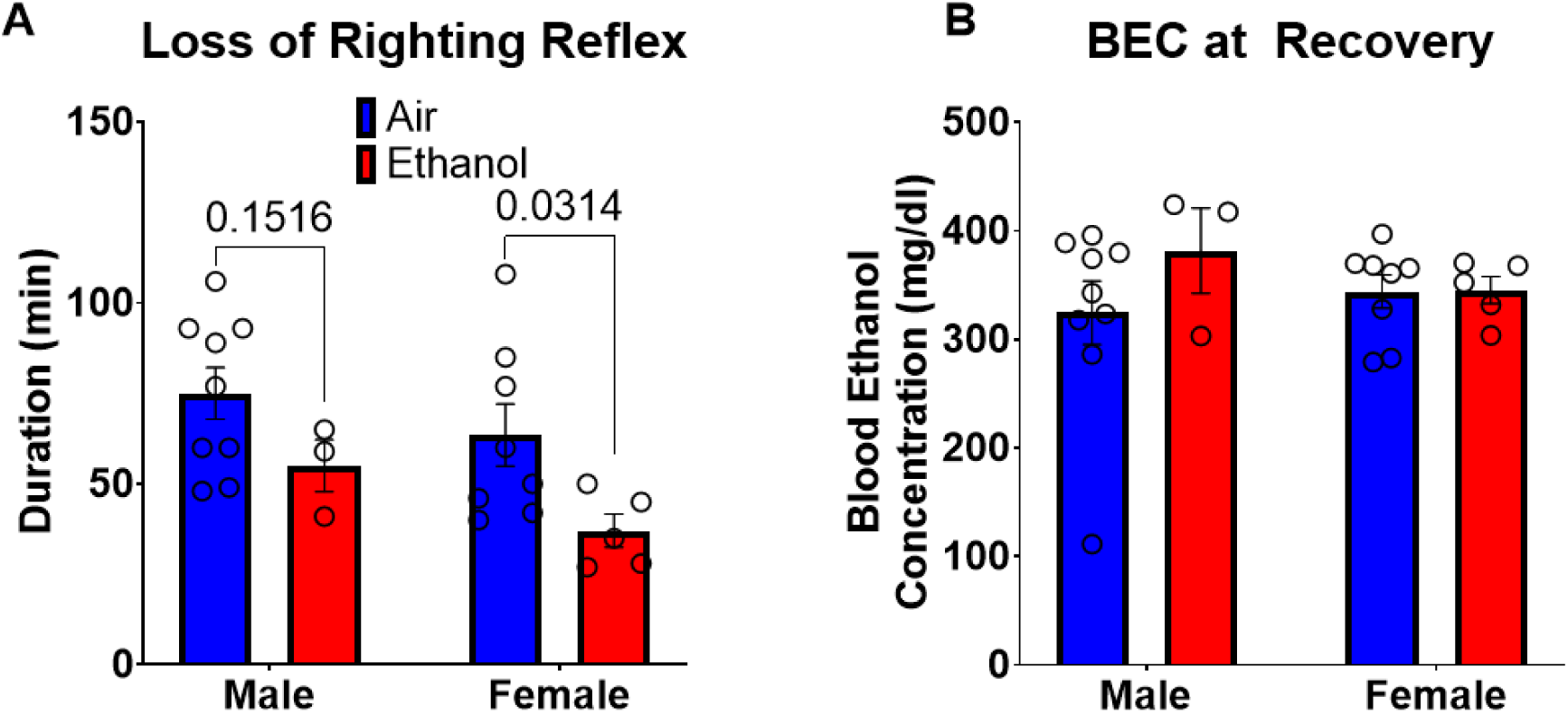
Loss of Righting Reflex. (A) Duration of loss of righting reflex in minutes. (B) Blood ethanol concentration at recovery of loss of righting reflex in mg/dl. All data are shown as mean 土 SEM.

### 3.5 Loss of Righting Reflex

On PND 116, all mice were tested for duration of LORR and their blood ethanol concentration was assessed upon recovery (**Figure 10**). Overall, ethanol-exposed mice recovered their righting reflex more quickly than air-exposed mice [Exposure (F_1, 21_ = 6.91, p < 0.02)], an effect that was largely driven by the female mice (**Figure 10A)**. However, there were no differences in BEC between groups regardless of adolescent AIR or AIE exposure (**Figure 10B**). It is important to note that only 3 out of 10 Male-AIE mice lost their righting reflex compared to 9 out of 10 Male-AIR mice lost their righting reflex. Only 5 out of 10 Female-AIE mice lost their righting reflex compared to 8 out of 10 Female-AIR mice.

## 4. Discussion

There is a growing body of research that links excessive alcohol consumption to changes in the gut microbiome, which can directly contribute to health issues such as alcohol use disorder and alcohol-associated liver disease (Wolstenholme et al. 2024; X. Wang et al. 2023). In addition, neuropsychiatric disorders can be caused by disruptions in the gut-brain axis (Wolstenholme et al. 2024). Metabolomics is also a newer methodological approach to serve as a measure identification of biomarkers in AUD and many other neuropsychiatric disorders (Harada et al. 2016). Specifically, cellular processes impacted by environmental stressors, such as sustained high levels of alcohol exposure, can be identified through metabolomic profiling. This includes noting alterations in amino acid, lipid, and carbohydrate metabolism (Harada et al. 2016; Harrigan, Maguire, and Boros 2008).

Chronic alcohol exposure can disrupt the gut-liver and gut-brain axes by altering the complex relationship between the gut and the liver through various mechanisms (Wolstenholme et al. 2024). Gut dysbiosis and impaired gut-liver metabolism resulting from microbiome changes are linked to the advancement of alcohol-induced liver disease (Ganesan et al. 2024). Human studies that compare alcohol users and nonusers have found that alcohol consumption alters the metabolome profiles of lipids and amino acids, which are linked to energy metabolism (Voutilainen and Kärkkäinen 2019; Clark et al. 2025). A rat model of AUD showed changes in lipid, amino acid, nucleotide, and carbohydrate metabolism pathways in fecal samples (X. Wang et al. 2023).

Ethanol exposure is indicated by the presence of two metabolites, lipoprotein and glucose (Du et al. 2020; Li et al. 2018; X. Wang et al. 2023). Binge ethanol exposure can hinder the molecular transport of albumin, transferrin, and glycoproteins, and lead to a decrease in protein synthesis, such as glycoprotein synthesis (Forman 1988). The decreased levels of amino acids found in the serum samples from our study support this conclusion. Oxidative stress, alterations in amino acid, and energy metabolism in animal models have been linked with models of anxiety-like behavior (Humer, Probst, and Pieh 2020; Humer, Pieh, and Probst 2020). Our findings align with previous research, as we also observed alterations in numerous amino acid metabolites including leucine, isoleucine, tryptophan, tyrosine, threonine, and methionine levels.

The open field test is a common measure of anxiety-like behavior in mice. This test utilizes the center zone as a measure of thigmotaxis; due to their natural aversion to bright open spaces, mice that display increased anxiety tend to avoid the center (Bolivar, Cook, and Flaherty 2000; Bailey and Crawley 2009; Lipkind et al. 2004; Simon, Dupuis, and Costentin 1994; Walz, Mühlberger, and Pauli 2016). Our research demonstrated that adult male mice exposed to ethanol during adolescence spent more time in the center zone and reared more than air-exposed males, exhibiting reduced anxiety-like behavior. The reasons behind this sex difference are unclear; however, using center zone data as a measure for ‘anxiety-like’ behavior, we can interpret these findings as suggesting that male mice exposed to ethanol during adolescence exhibit a reduced ‘anxiety-like’ behavioral phenotype compared to air-exposed male mice. These data indicate long-term dynamic relationships between adolescence exposure in the context of sex and anxiety (Peeters et al. 2024).

A human study conducted by Leclercq et al. investigated the relationship between affective behaviors (anxiety, depression, and alcohol craving) and metabolomic changes in patients with and without AUD (Leclercq et al. 2024). The researchers found that the metabolome profiles of patients with AUD differed from those without AUD. Additionally, in some cases, abstinence led to a return to levels seen in healthy controls (Leclercq et al. 2024). Recent work has assessed similar changes induced by alcohol dependence in behavioral changes and changes in fecal and serum metabolite profiles after chronic intermittent two bottle choice drinking in adult Sprague Dawley rats. Specifically, these rats were tested on the 28th day of intermittent ethanol exposure (X. Wang et al. 2023). Similar to our present findings,Wang and colleagues showed changes in anxiety-like behavior in the open field test and elevated plus maze and separations in metabolite profiles in serum samples (X. Wang et al. 2023). Our work showed moderate changes in anxiety-like behavior and correlations between metabolites in serum, fecal, and liver samples. Our work differed from previous research in several key ways. First, it included both male and female subjects. Second, it involved multiple fecal sample collections at different time points throughout the ethanol withdrawal process. Third, it examined changes in liver metabolites that occurred after acute intermittent ethanol exposure.

Costanzo and colleagues examined sex differences in plasma metabolomics in humans and found that men tended to have higher levels of amino acids than women. These amino acids included phenylalanine, glutamate, glutamine, kynurenine, methionine, proline, and tyrosine. Additionally, men had higher levels of the branched-chain amino acids (BCAA) valine, leucine, and isoleucine (Costanzo et al. 2022). In human men, higher alcohol consumption was associated with levels of serum threonine, glutamine, and guanidinosuccinate (Harada et al. 2016). In serum samples, lipid, amino acid, and carbohydrate metabolism were differently affected by intermittent ethanol exposure in the adult male rats (X. Wang et al. 2023). Similar to recent work, we also found changes in metabolites in the ethanol-exposed group compared to the air-exposed group in serum and fecal samples, including proline, glycine, serine, tryptophan, lysine, and threonine. These metabolites are linked to pathways involved in oxidative stress and immune functions. The AIE-induced differences in male and female mice observed in the current study exhibited a very similar metabolite profile to those seen in previous research.

Changes in epigenetic modifications, addiction, and reward networks in the brain are associated with alterations in specific metabolites, including butyrate (Wolstenholme et al. 2024). Other work in humans showed specific changes in the gut microbiome associated with AUD and alcoholic liver disease, including a decrease in butyrate-producing bacteria and an increase in endotoxin-producing bacteria (Litwinowicz and Gamian 2023). In our current work, butyrate was associated with anxiety-like behavior in the short-term withdrawal fecal samples in male mice. Similar to our work, there were some distinct metabolites altered in serum compared to fecal samples after ethanol exposure (X. Wang et al. 2023).

Perturbations in metabolic processes can induce a greater susceptibility to developing AUD and associated disorders after prolonged use (Harrigan, Maguire, and Boros 2008). Many of the deleterious effects of heavy alcohol use may be secondary to alcohol’s impact on liver function (Harrigan, Maguire, and Boros 2008). To date there is limited research examining changes in liver metabolomic profiles in rodent models of heavy alcohol exposure. To our knowledge this is the first study to date to determine long-term changes in liver metabolic profiles following adolescent ethanol exposure in mice.

A potential limitation of this study was the use of NMR-based analysis, rather than LC/MS/MS techniques, which may have limited the identification of differentially expressed metabolites between groups (Voutilainen and Kärkkäinen 2019; Yang et al. 2024).The different collection times and potential compensatory changes between sample types made it difficult to identify which metabolites were altered across withdrawal periods (Voutilainen and Kärkkäinen 2019). Our data presented here is a semi-cross-sectional analysis of alcohol-induced metabolite profiles using different sample types at different withdrawal time points after AIE exposure in male and female mice.

The metabolome provides insight into functional changes and disease progression resulting from heavy alcohol use. Compared to other ‘omics’ approaches, the metabolome offers a deeper understanding of the overall functional alterations caused by metabolic processes (Voutilainen and Kärkkäinen 2019). Several types of samples, including serum, urine, and fecal samples, can be used to test for metabolomic changes following heavy alcohol exposure including in humans and preclinical models (Harada et al. 2016; Harrigan, Maguire, and Boros 2008; X. Wang et al. 2023; Yang et al. 2024). Targeted and nontargeted metabolomic approaches can be used to determine changes in metabolites that are altered after heavy alcohol exposure, with both having advantages and disadvantages (Harrigan, Maguire, and Boros 2008). We employed a nontargeted metabolomic approach to comprehensively characterize the alterations caused by heavy alcohol consumption during adolescence.This initial research aimed to establish a proof-of-concept model by analyzing how ethanol exposure during adolescence affects metabolic profiles in male and female mice. This was achieved by examining various sample types, including serum, fecal, and liver samples.

Metabolomics approaches can serve to help in treating afflicted individuals of AUD with a more personalized or tailored approach (Harrigan, Maguire, and Boros 2008). Goffredo and colleagues found that obese adolescents with NAFLD have higher plasma levels of certain amino acids (valine, isoleucine, tryptophan, and lysine) compared to those without NAFLD (Goffredo et al. 2017). These elevated amino acid levels were associated with decreased insulin sensitivity and predicted an increase in hepatic fat content over time, independent of obesity and insulin resistance. These data together with those in the present work examining changes in affective behaviors, metabolite profiles in several different sample types following AIE, and incorporating sex as a variable can inform future work aimed at identifying specific metabolic biomarkers important for AUD progression.

## Data Availability Statement

The data that support the findings of this study are available from the corresponding author upon reasonable request.

## Author Contributions

MJS, AK, NIH, RCW, CEE, DH, MEF, BW, AMD collected the data. MJS, AK, MP, JAB, TD, RC, BW and AMD drafted the manuscript. MJS, BW, and AMD designed the experiments. MJS, AK, BW and AMD analyzed the data. MJS, AK, NIH, RCW, JAB, CEE, TD, RC, BW and AMD edited the manuscript.

## Other Acknowledgements

The authors would like to thank Destiny M. Belton, Taylor Costen, Victoria Robinson, Frederick Feely, and Pragya Nepal for their expert technical assistance.

