## Supplemental Data for "Adolescent intermittent ethanol exposure induces sex-specific and time-dependent changes in affective behaviors and metabolomic profiles"

Supplemental Data: *Metabolomics: Activity relationship with serum, fecal, and liver samples*

### 1.1 Serum samples

The correlation relationship of serum metabolites with the activities data were analyzed using the Pearson correlation relationship study. They are shown in **Table 6** below. The serum samples were collected on PND 43.

### Table S1: Correlation between behavior measures and serum metabolites

| | Air Male | | EtOH Male | | Air Female | | EtOH Female | |
|---|---|---|---|---|---|---|---|---|
| **Alanine** | | | TDM<br>Center Zone Entries<br>Center Zone TDM | r = 0.83, p = 0.041<br>r = 0.89, p = 0.01<br>r = 0.86, p = 0.02 | Center Zone Time | r = -0.87, p = 0.02 | | |
| **Citrate** | | | Center Zone Time | r = -0.83, p = 0.03 | | | | |
| **Glucose** | | | TDM<br>Center Zone TIme<br>Center Zone Entries<br>Center Zone TDM | r = -0.95, p = 0.003<br>r = -0.95, p = 0.003<br>r = -0.90, p = 0.01<br>r = 0.98, p = 0.0005 | | | | |
| **Glutamate** | | | Center Zone Time | r = -0.95, p = 0.002 | Center Zone Time | r = -0.83, p = 0.04 | | |
| **Glutamine** | LDT Time | r = -0.83, p = 0.03 | Center Zone Time | r = -0.92, p = 0.009 | | | Rearing | r = 0.97, p = 0.001 |
| **Isoleucine** | | | LDT Time | r = 0.85, p = 0.02 | | | TDM<br>Center Zone Entries<br>Center Zone TDM | r = 0.83, p = 0.03<br>r = 0.91, p = 0.01<br>r = 0.86, p = 0.02 |
| **Lactate** | | | | | Center Zone Time<br>Center Zone Entries<br>Center Zone TDM | r = 0.87, p = 0.02<br>r = 0.87, p = 0.02<br>r = 0.89, p = 0.01 | | |
| **Leucine** | | | Center Zone TDM | r = 0.84, p = 0.03 | LDT Time | r = -0.92, p = 0.008 | | |
| **Lipoprotein** | | | TDM<br>Center Zone Time<br>Center Zone Entries<br>Center Zone TDM | r = 0.85, p = 0.03<br>r = 0.98, p = 0.006<br>r = 0.82, p = 0.046<br>r = 0.88, p = 0.01 | | | | |

| | | | | | | |
|---|---|---|---|---|---|---|
| **Phenylalanine** | | | Center Zone Time<br>Center Zone TDM | r = -0.90, p = 0.01<br>r = -0.83, p = 0.03 | | |
| **Proline** | TDM<br><br>Center Zone Entries<br>Center Zone TDM | r = 0.83, p = 0.04<br>r = 0.83, p = 0.041<br>r = 0.83, p = 0.042 | Center Zone Time<br>Center Zone TDM | r = -0.87, p = 0.02<br>r = 0.84, p = 0.03 | TDM<br><br>Rearing<br><br>Center Zone Entries<br>Center Zone TDM | r = -0.85, p = 0.03<br>r = -0.96, p = 0.002<br>r = -0.89, p = 0.01<br><br>r = -0.87, p = 0.02 |
| **Succinate** | | | Center Zone Time<br>Center Zone TDM | r = -0.94, p = 0.004<br>r = -0.84, p = 0.03 | | |
| **sn-Glycero-3-phosphocholine** | | | TDM<br><br>Center Zone TDM | r = 0.91, p = 0.01<br>r = 0.89, p = 0.01 | | |
| **Taurine** | | | Center Zone Time | r = -0.85, p = 0.03 | | |
| **Tyrosine** | | | LDT Time | r = -0.83, p = 0.03 | | |

#### 1.2 Fecal samples short-term withdrawal

The correlation relationship of early withdrawal fecal metabolites with the activities data were analyzed using the Pearson correlation relationship study. They are shown in **Table 7** below. The fecal samples were collected between PND 49-53.

### Table S2: Correlation between behavior measures and short-term withdrawal fecal metabolites

| | Air Male | | EtOH Male | | Air Female | | EtOH Female | |
|---|---|---|---|---|---|---|---|---|
| **1,3-Dihydroxyacetone** | TDM<br>Center Zone Time<br>Center Zone Entries<br>Center Zone TDM | r = 0.94; p = 0.004<br>r = 0.85; p = 0.03<br>r = 0.93; p = 0.005<br>r = 0.93; p = 0.007 | TDM<br>Center zone TDM | r = -0.87; p = 0.02<br>r = -0.82; p = 0.048 | LDT Time | r = 0.83; p = 0.041 | | |
| **3-Methyl-2-oxovalerate** | TDM<br>Center Zone Time<br>Center Zone Entries<br>Center Zone TDM | r = -0.93; p = 0.007<br>r = -0.81; p = 0.049<br>r = 0.95; p = 0.003<br>r = -0.91; p = 0.01 | | | | | | |
| **Acetoin** | | | | | LDT Time | r = 0.81; p = 0.049 | | |
| **Alanine** | | | | | LDT Time | r = -0.94; p = 0.005 | | |
| **Asparagine** | | | Center zone TDM<br>LDT Time | r = -0.82; p = 0.045<br>r = -0.92; p = 0.008 | | | | |
| **Aspartate** | Center zone time | r = -0.87; p = 0.02 | TDM<br>Center zone time<br>Center zone TDM | r = -0.88; p = 0.02<br>r = -0.83; p = 0.04<br>r = -0.86; p = 0.02 | | | | |
| **Butyrate** | TDM<br>Center zone time<br>Center Zone Entries<br>Center Zone | r = -0.93; p = 0.006<br>r = 0.83; p = 0.04<br>r = -0.82; p = 0.004<br>r = 0.91; p = 0.01 | LDT Time | r = 0.84; p = 0.03 | | | | |
| **Formate** | | | Center zone time | r = -0.82; p = 0.046 | LDT Time | r = 0.92; p = 0.009 | Center zone time<br>Center zone entries<br>Center zone TDM | r = 0.84; p = 0.03<br>r = 0.91; p = 0.01<br>r = 0.89; p = 0.01 |
| **Fumarat** | Center Zone | r = -0.83; p | | | | | | |

| | | | | | | | | |
|---|---|---|---|---|---|---|---|---|
| **e** | Entries | = 0.04 | | | | | | |
| **Glucose** | | | TDM | R = 0.90; p = 0.01 | | | | |
| **Glutamate** | TDM | r = -0.83; p = 0.04 | | | | | | |
| **Glycocholate** | TDM<br>Center Zone Entries<br>Center Zone TDM | r = -0.84; p = 0.04<br>r = -0.84; p = 0.036<br>r = -0.81; p = 0.048 | | | | | Center zone time<br>Center zone TDM | r = 0.88; p = 0.02<br>r = 0.86; p = 0.02 |
| **Hypoxanthine** | TDM<br>Center Zone TDM | r = -0.81; p = 0.049<br>r = 0.82; p = 0.04 | | | | | | |
| **Lactate** | LDT Time | r = -0.83; p = 0.04 | TDM<br>Center zone TDM<br>LDT Time | r = -0.85; p = 0.03<br>r = -0.89; p = 0.01<br>r = -0.84; p = 0.03 | LDT Time | r = 0.88; p = 0.02 | | |
| **Leucine** | | | | | | | Center zone time | r = 0.81; p = 0.049 |
| **Nicotinate** | TDM<br>Center Zone Time<br>Center Zone TDM | r = 0.99; p = 0.0001<br>r = 0.89; p = 0.017<br>r = 0.99; p = 0.0002 | | | LDT Time | r = 0.84; p = 0.03 | Center zone time<br>Center zone TDM | r = -0.84; p = 0.03<br>r = -0.88; p = 0.01 |
| **Phenylalanine** | TDM<br>Center Zone Time<br>Center Zone Entries<br>Center Zone TDM1 | r = 0.86; p = 0.03<br>r = 0.97; p = 0.001<br>r = 0.87; p = 0.02<br>r = 0.90; p = 0.01 | | | | | Center zone time<br>Center zone TDM | r = -0.82; p = 0.044<br>r = -0.84; p = 0.03 |
| **Saccharopine** | TDM | r = -0.82; p = 0.048 | | | | | | |
| **Succinate** | Rearing | r = 0.93; p = 0.007 | | | | | | |
| **Taurine** | | | | | | | TDM | r = 0.85; p = 0.03 |
| **Threonine** | | | TDM<br>Center zone time<br>Center | r = -0.99; p = 0.0002<br>r = -0.84; p = 0.03<br>r = -0.93; p | LDT Time | r = 0.86; p = 0.03 | | |

| | | | | | | | | |
|---|---|---|---|---|---|---|---|---|
| | | | zone entries<br>Center zone TDM | = 0.006<br><br>r = -0.96; p = 0.002 | | | | |
| **Total Bile Acid** | | | Center zone time | r = -0.86; p = 0.02 | | | | |
| **Trimethylamine** | | | Rearing | r = 0.82; p = 0.045 | | | | |
| **Tryptophan** | | | | | LDT Time | r = 0.91; p = 0.01 | Rearing | r = -0.98; p = 0.0007 |
| **Tyrosine** | | | | | | | TDM | r = -0.88; p = 0.01 |

### 1.3 Fecal samples long-term withdrawal

The correlation relationship of late withdrawal fecal metabolites with the activities data were analyzed using the Pearson correlation relationship study. They are shown in **Table 8** below. The fecal samples were collected between PND 91-95.

**Table S3: Correlation between behavior measures and long-term withdrawal fecal metabolites**

| | Air Male | | EtOH Male | | Air Female | | EtOH Female | |
|---|---|---|---|---|---|---|---|---|
| **1,3-Dihydroxyaceton** | TDM<br><br>Center Zone TDM | r = -0.93; p = 0.01<br>r = -0.93; p = 0.01 | | | | | | |
| **Asparagine** | | | | | Rearing | r = 0.82; p = 0.04 | | |
| **Aspartate** | LDT Time | r = -0.90; p = 0.03 | | | | | | |
| **Choline** | | | Center Zone Time | r = 0.92; p = 0.02 | | | | |
| **Formate** | Center Zone Time<br>LDT Time | r = 0.93; p = 0.02<br>r = 0.95; p = 0.01 | | | | | | |
| **Glucose** | Center Zone Time<br>LDT Time | r = -0.92; p = 0.02<br>r = -0.93; p = 0.01 | LDT Time | r = 0.89; p = 0.04 | | | | |
| **Glutamine** | Center Zone Time | r = 0.99; p = 0.001 | | | | | | |
| **Hypoxanthine** | Center Zone Entries | r = 0.93; p = 0.01 | | | | | | |
| **Nicotinate** | | | | | | | Center Zone TIme | r = 0.82; p = 0.04 |
| **Phenylalanine** | Rearing<br><br>LDT Time | r = 0.98; p = 0.002<br>r = -0.90; p = 0.03 | | | Center Zone Time<br>Center Zone Entries | r = 0.94; p = 0.004<br>r = 0.88; p = 0.02 | | |
| **Proprionate** | | | Rearing | r = -0.94; p = 0.01 | | | LDT Time | r = 0.81; p = 0.047 |

| | | | | | | | | |
|---|---|---|---|---|---|---|---|---|
| **Saccharopine** | | | | | | | LDT Time | r = 0.93; p = 0.006 |
| **Succinate** | LDT Time | r = 0.95; p = 0.01 | | | | | | |
| **Taurine** | | | | | | | Rearing | r = 0.89; p = 0.01 |
| **Threonine** | | | LDT Time | r = -0.91; p = 0.02 | | | LDT Time | r = 0.81; p = 0.047 |
| **Trimethylamine** | LDT Time | r = 0.96; p = 0.007 | | | TDM<br>Center Zone Entries<br>Center Zone TDM | r = 0.87; p = 0.02<br>r = 0.87; p = 0.02<br>r = 0.87; p = 0.02 | | |
| **Typtophan** | | | | | TDM<br>Center Zone Entries<br>Center Zone TDM | r = 0.81; p = 0.05<br>r = 0.95; p = 0.003<br>r = 0.81; p = 0.05 | Center Zone Time | r = 0.85; p = 0.03 |
| **Tyrosine** | Rearing<br>LDT Time | r = 0.94; p = 0.01<br>r = 0.96; p = 0.008 | | | Rearing<br>Center Zone Entries | r = 0.87; p = 0.02<br>r = 0.92; p = 0.009 | | |
| **Xanthine** | Center Zone Time | r = 0.90; p = 0.03 | | | Center Zone TIme | r = 0.81; p = 0.04 | | |

### 1.4 Liver samples long-term withdrawal

The correlation relationship of liver metabolites with the activities data from the last behavioral trial were analyzed using the Pearson correlation relationship study. They are shown in **Table 9** below. The liver samples were collected on PND 119.

**Table S4: Correlation between behavior measures and liver metabolites**

| | Air Male | | EtOH Male | | Air Female | | EtOH Female |
|---|---|---|---|---|---|---|---|
| **Acetate** | Rearing | r = 0.82; p = 0.041 | | | | | |
| **Betaine** | Center Zone TDM | r = -0.86; p = 0.02 | | | | | |
| **Choline** | Rearing | r = 0.93; p = 0.007 | | | | | |
| **Creatinine** | | | Center Zone Entries<br>Center Zone TDM | r = 0.81; p = 0.02<br>r = 0.78; p = 0.03 | | | |
| **Glutamate** | Rearing | r = 0.82; p = 0.043 | Center Zone Entries<br>Center Zone TDM | r = 0.79; p = 0.03<br>r = 0.78; p = 0.03 | | | |
| **Glycine** | | | | | LDT Time | r = 0.89; p = 0.03 | |
| **Inosine** | | | Center Zone Entries<br>Center Zone TDM | r = -0.88; p = 0.008<br>r = -0.88; p = 0.008 | | | |
| **Isoleucine** | Rearing | r = 0.84; p = 0.03 | | | | | |
| **Isocitrate** | | | LDT Time | r = 0.91; p = 0.004 | | | |
| **Lactate** | Center Zone TDM | r = 0.82; p = 0.043 | | | | | |
| **Leucine** | Rearing | r = 0.94; p = 0.004 | | | | | |

| | | | | | | |
|---|---|---|---|---|---|---|
| **O-phosphocholine** | | | Rearing | r = -0.83; p = 0.01 | | |
| **Phenylalanine** | | | TDM<br><br>Center Zone Entries | r = 0.78; p = 0.03<br>r = 0.75; p = 0.048 | | |
| **Pyruvate** | | | Center Zone Entries | r = 0.78; p = 0.03 | | |
| **Taurine** | Center Zone Time | r = 0.92; p = 0.008 | | | Rearing | r = -0.82; p = 0.041 |
| **Tyrosine** | | | Center Zone Entries<br>Center Zone TDM | r = 0.80; p = 0.02<br><br>r = 0.78; p = 0.03 | | |
| **UDP-galactose** | | | LDT Time | r = 0.77; p = 0.04 | | |
